# Deconstructing cell-free extract preparation for *in vitro* activation of transcriptional genetic circuitry

**DOI:** 10.1101/411785

**Authors:** Adam D. Silverman, Nancy Kelley-Loughnane, Julius B. Lucks, Michael C. Jewett

## Abstract

Recent advances in cell-free gene expression (CFE) systems have enabled their use for a host of synthetic biology applications, particularly for rapid prototyping of genetic circuits designed as biosensors. Despite the proliferation of cell-free protein synthesis platforms, the large number of currently existing protocols for making CFE extracts muddles the collective understanding of how the method by which an extract is prepared affects its functionality. Specifically, a key goal toward developing cell-free biosensors based on native genetic regulators is activating the transcriptional machinery present in bacterial extracts for protein synthesis. However, protein yields from genes transcribed *in vitro* by the native *Escherichia coli* RNA polymerase are quite low in conventional crude extracts originally optimized for expression by the bacteriophage transcriptional machinery. Here, we show that cell-free expression of genes under bacterial σ^70^ promoters is constrained by the rate of transcription in crude extracts and that processing the extract with a ribosomal run-off reaction and subsequent dialysis can alleviate this constraint. Surprisingly, these processing steps only enhance protein synthesis in genes under native regulation, indicating that the translation rate is unaffected. We further investigate the role of other common process variants on extract performance and demonstrate that bacterial transcription is inhibited by including glucose in the growth culture, but is unaffected by flash-freezing the cell pellet prior to lysis. Our final streamlined protocol for preparing extract by sonication generates extract that facilitates expression from a diverse set of sensing modalities including protein and RNA regulators. We anticipate that this work will clarify the methodology for generating CFE extracts that are active for biosensing and will encourage the further proliferation of cell-free gene expression technology for new applications.

## INTRODUCTION

Cell-free systems are emerging as a prominent platform for use in synthetic biology [1, 2]. By mixing clarified cellular extracts, exogenously supplied energy sources and cofactors, and DNA-encoded genetic instructions, cell-free protein synthesis supports the basic processes of gene expression and metabolism in a convenient and engineerable *in vitro* reaction environment. Because the open environment enables flexibility for optimizing extract and reaction conditions and is amenable to high-throughput automation [3], cell-free gene expression (CFE) technology has found great utility in a wide range of contexts. Since their first application in deciphering the genetic code [4, 5], cell-free systems have been successfully applied for the bulk production of model [6-9] and therapeutic proteins [10-15]. Beyond just protein synthesis, though, CFE technologies have evolved more generally to enable complex and diverse functions, including prototyping cellular metabolism [16-18] and glycosylation [19-21], expressing minimal synthetic cells, virus-like particles, and bacteriophages [7, 22-26], portable on-demand manufacturing of pharmaceuticals [27, 28], incorporation of nonstandard amino acids within proteins [29-33], prototyping of genetic circuitry [34-36], and sensing viral RNAs and small molecules through rapid, low-cost, and field-deployable molecular diagnostics [37-42]. Most progress has occurred in CFE systems generated from *Escherichia coli* strains engineered for protein production, largely due to the bacterium’s well-characterized genetics and metabolism [1]. However, there has been recent progress in adapting CFE protocols to make lysates from eukaryotic and non-model organisms, including yeast [43, 44], Gram-positive bacteria [45, 46], plants [47, 48], and mammalian cells [49-51]. CFE technology is therefore now at the point of expanding beyond specialist laboratories and becoming a major toolbox throughout synthetic biology research, application, and recently, education [34, 52, 53].

A cell-free gene expression reaction is composed of three or four main components that enable *in vitro* gene expression and metabolism: the clarified cellular lysate (or “extract”) that contains the requisite cellular machinery for protein synthesis; a buffered mixture of phosphorylated energy substrates, NTPs, amino acids, salts, and other required cellular cofactors; the DNA templates that define the genetic program to be executed in the reaction; and any other exogenous cofactors, substrates, or inducers required for the reactions. Of these, the extract is the most labor-intensive component to prepare, requiring precise control over cell culture fermentation, lysis, and post-lysis separation of unwanted cellular debris from the transcriptional and translational machinery that must remain behind in the final extract. Recent work has focused on optimizing performance of and expanding access to CFE technology by simplifying extract preparation protocols, including replacing lysis by homogenization with cheaper methods like sonication [54, 55], bead-beating [54, 56], enzymatic lysis [57], or flash-freezing [58], as well as reducing centrifugation intensity to speeds accessible on conventional benchtop instruments [59]. While these efforts have made great strides, no standard method yet exists for preparing highly active *E. coli* extracts, creating a barrier against the wider adoption of CFE systems in the field. Moreover, as a result of protocol and performance inconsistencies between research groups, many labs instead opt to use chemically defined, bottom-up reconstituted cell-free gene expression systems such as the “purified recombinant elements” (PURE) system [37, 60, 61]. Although reconstituted *in vitro* protein synthesis platforms are powerful, their cost can be prohibitive, and they also lack the flexibility for strain engineering and cofactor and energy regeneration afforded by cellular extracts. Thus, it is still an important goal to develop a standardized method for extract preparation that would enable inexpensive, robust and engineerable crude extract CFE systems that are useful for a range of applications.

One of the major difficulties in standardizing CFE extract preparation is deconstructing the effects of different protocol variations on extract performance. Because the physiochemical environment of the extract is still poorly understood, little is known about how specific variations in the protocol used to prepare extract impact its utility for different applications (*e.g.*, high protein expression titers versus active genetic circuitry). In this work, we set out to characterize one such performance inconsistency: the functional inactivity of simple genetic programs using native regulatory elements in an extract that had been previously optimized for bulk protein production. Specifically, we discovered that extracts optimized to yield high protein titers above 1000 ng/μL using T7 bacteriophage promoters to drive model reporter protein expression had 15-fold lower protein expression under the control of native *E. coli* σ^70^ promoters. Because transcriptional regulation of the native *E. coli* polymerase is crucial for many applications of CFE systems, we aimed to uncover which aspect of the extract preparation process caused this discrepancy, toward the goal of generating an improved CFE platform that supports gene expression from native *E. coli* regulators.

Here, we demonstrate that the transcriptional limitations from *E. coli* regulatory elements are removed by the addition of specific post-lysis processing steps in the preparation of crude extracts for CFE. Specifically, we find that ribosomal run-off and dialysis steps are critical for recovering transcriptional activity from *E. coli* σ^70^ promoters. We also investigate the effects of other common extract preparation steps such as flash freezing and evaluate their impact on final extract performance. Combining these features, we create a modified cell-free extract preparation framework that can be done in a single twelve-hour day or split across two shorter days and demonstrate its batch-batch reproducibility. Overall, this system improves the cell-free gene expression yield from bacterial σ^70^ promoters by five-fold, speeds up the signal response time by three-fold, and can be used to activate of a wide array of synthetic genetic regulators and circuits, including RNA transcriptional activators and repressors, protein transcription factors, CRISPR-Cas9 repression, toehold switches, and riboswitches. Overall, this work serves to illuminate the “black box” of CFE and make cell-free technology more accessible, particularly for the numerous applications that require cell-free expression from the bacterial transcriptional machinery. We anticipate that this work will have implications in cell-free synthetic biology and molecular diagnostics.

## Results and discussion

### Cell-free gene expression from native bacterial promoters is severely limited

A rapidly growing application area for bacterial cell-free systems is to use them for prototyping genetic elements (*e.g.*, promoters, terminators) and circuits for function in living hosts [34]. We therefore first set out to assess the ability to activate native transcriptional machinery in the sonication-based CFE platform developed and optimized for protein production by Kwon and Jewett [55]. We began by preparing cell free extracts from the Rosetta2 (DE3) pLysS strain of *Escherichia coli*, a derivative of *E. coli* BL21 supplemented with a plasmid encoding “rare” bacterial tRNAs to facilitate enhanced translation, making it optimal for recombinant protein production. After extracts were prepared following our published sonication-based protocol [55] (Figure 1A, “No processing”, gray line), we carried out cell-free gene expression of the model reporter superfolder green fluorescent protein (sfGFP). The sfGFP expression cassette contained a T7 bacteriophage promoter followed by the sfGFP coding sequence and T7 transcriptional terminator. A batch-mode transcription-translation reaction was carried out by adding the expression cassette template plasmid DNA to a mixture containing T7 RNA polymerase (RNAP), cell extract, and essential substrates (*e.g.*, amino acids, nucleotides, energy substrates, cofactors, and salts) to a final volume of 15 μL and incubating this mixture at 30^°^C for 15 hours. sfGFP expression was quantified at the end of the reaction by measuring fluorescence (485 nm excitation, 520 nm emission) using a plate reader, which was then compared to a standard curve to derive protein concentrations. Consistent with previous work, our extract yielded greater than 1000 ng/μL of sfGFP, confirming that it was active for bulk protein production.

**Figure 1.**
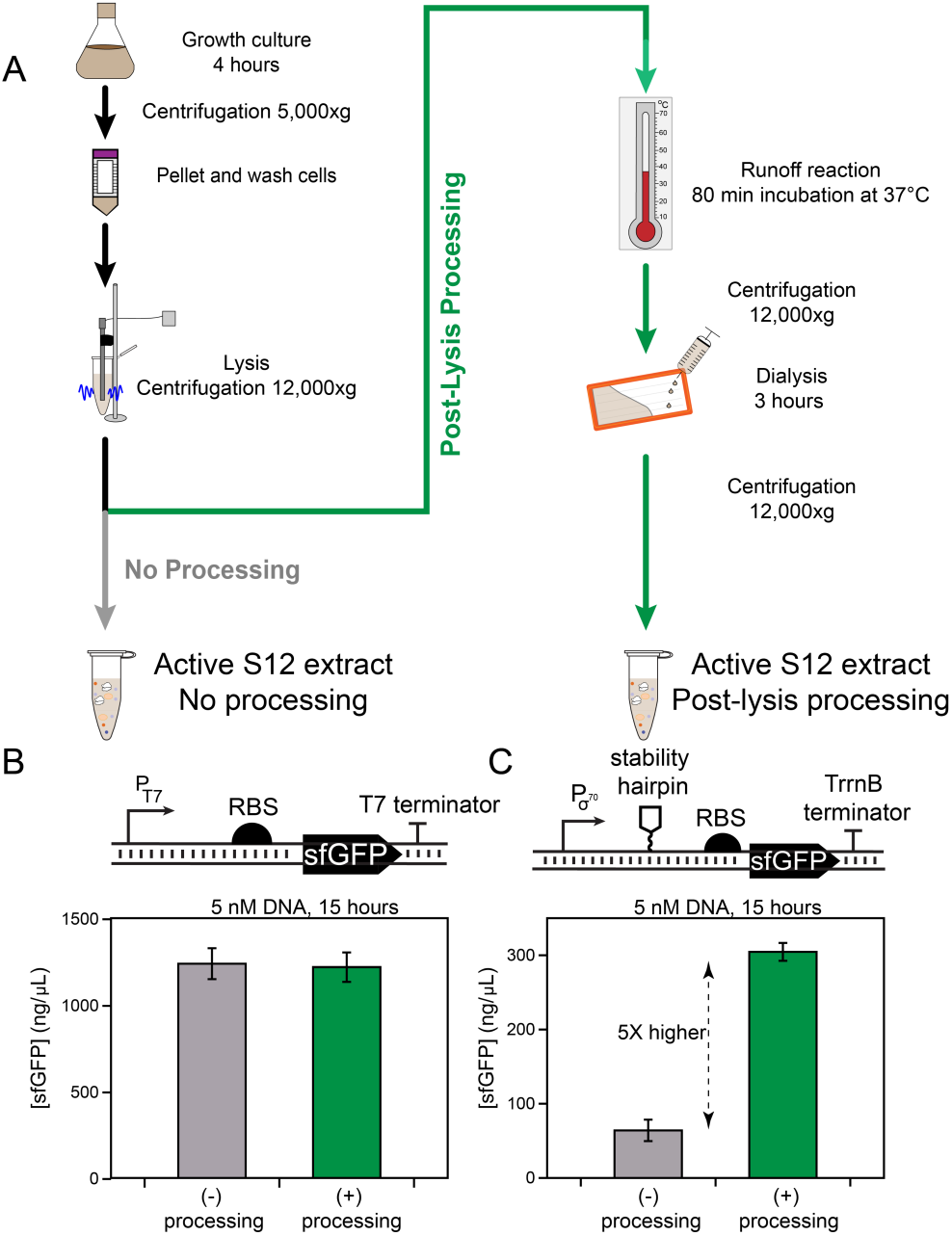
The impact of different extract preparation procedures on cell-free gene expression (CFE) from different promoter systems. (a) Schematic for extract preparation. S12 extract preparation begins with culturing an *E. coli* strain (Rosetta2 pLysS) in enriched 2X YT+P media (see Methods) to optical density 3.0. Cultures are then processed through centrifugation, resuspension of the cell pellet, and lysis by physical disruption or sonication. The lysate is clarified by centrifugation at 12,000xg and either directly aliquoted (following [55], gray line), or post-lysis processed, consisting of an incubation at 37^°^C for 80 minutes (ribosomal run-off reaction) followed by centrifugation, dialysis for three hours through a 10 kDa molecular-weight cut-off cassette, and one final centrifugation step (this work, green line). Combined transcription and translation activity differs based on extract preparation procedure and the use of endogenous or exogenous (bacteriophage) promoter systems. (b) Extract preparations that did not include post-lysis processing steps yielded protein titers of ∼1250 ng/μL when the sfGFP reporter is expressed from a T7 promoter (P_T7_) using exogenously added T7 RNA polymerase, independent of extract preparation steps. (c) Extracts prepared without post-lysis processing showed poor yield (∼50 ng/μL) when under the control of the consensus *E. coli* σ^70^ promoter (P_σ70_) and *E. coli* RNA polymerase supplied from the lysate. The addition of processing steps to the lysate outlined in green in (a) improved protein expression yields from the bacterial σ^70^ promoter by 5X without impacting yields from the T7 promoter system. Protein yields are from overnight (15-hour) in 15 μL reactions carried out in 2.0 mL tubes. Reporter constructs using the consensus *E. coli* σ^70^ promoter also encoded an RNA stability hairpin before the RBS and downstream coding sequence (Figure S1). Reactions were supplemented with 5 nM of a plasmid-based sfGFP expression cassette in 2.0 mL tubes, and protein yields were background-subtracted (i.e., no plasmid control) and measured by correlation to a known fluorescent standard. Error bars represent the standard deviation of the mean from three independent reactions.

After validating gene expression under control of T7 RNAP, we next sought to investigate production of sfGFP under control of a bacterial σ^70^ promoter and the *E. coli* RNAP already present in the extract. Previous work has shown that bacterial cell-free gene expression systems can support expression from an array of native prokaryotic transcriptional components [7, 62]. To verify this in our system, we constructed an sfGFP expression cassette containing the consensus *E. coli* σ^70^ promoter followed by the sfGFP coding sequence and the TrrnB transcriptional terminator from the *E. coli* rrnB ribosomal operon. To our surprise, we observed no expression of the reporter protein above background when this DNA construct was incubated with extract and transcription-translation reagents for 15 hours at 30^°^C (Supplementary Figure S1). Drawing inspiration from previous work that reported substantial improvements to cell-free expression productivity when strong secondary structure is added to the 5’ untranslated region of the reporter mRNA [62], we modified our expression construct to include the PHP14 stability hairpin [63] immediately upstream of the ribosome-binding site on our reporter (Refer to the Supplementary Plasmid Files for more detailed information on plasmid architecture). With this construct, we observed 64 ± 14 ng/μL of constitutive protein production in the same reaction conditions, which was about 20-fold lower than that observed from the T7 promoter (Figure 1B and 1C, gray bar). Since such low yields from cell-free expression would likely be insufficient for prototyping genetic circuits with naturally low ON states, we sought to optimize our extract preparation protocol to favor better cell-free productivity using the endogenous bacterial transcriptional machinery.

### Post-lysis processing significantly enhances CFE yield from native bacterial promoters

We next sought to uncover which steps in the extract preparation protocol were responsible for poor expression from bacterial promoters. CFE extract preparation consists of five key steps: pre-culture, growth culture, cell harvest, cell lysis, and post-lysis processing (Figure 1). Historically, a number of protocols have been developed for making productive extract, principally varying in the lysis and post-lysis steps (Table 1) [4, 5, 55, 56, 59]. Due to its great potential for scalability, equipment accessibility, and relatively low cost, we considered sonication as the preferable lysis method compared to physical lysis techniques such as homogenization or bead-beating. We therefore aimed to fix sonication as a design constraint while optimizing our extract preparation methods to enhance expression from the native transcription machinery.

**Table 1:** Outline of protocol differences between extract preparation methods. Italicized: variables examined in this study.

| **Extract Protocol** | **Zubay, 1973 [4]** | **Sun et. al., 2013 [56]** | **Kwon & Jewett, 2015 [55]** | **This work** |
| --- | --- | --- | --- | --- |
| Starter culture | One 16-hour culture | Two 8-hour cultures | One 16-hour culture | One 16-hour culture |
| *Culture media composition* | 2X YT | 2X YT + phosphate | 2X YT + phosphate + glucose | 2X YT + phosphate |
| Lysis method | French press | Bead-beating | Sonication | Sonication |
| Post-lysis centrifugation | 30,000xg for 30 minutes | 12,000xg for 10 minutes | 12,000xg for 10 minutes | 12,000xg for 10 minutes |
| *Run-off reaction* | 80 minutes | 80 minutes | Strain-dependent; none for BL21 | 80 minutes |
| *Dialysis* | 18 hours | 3 hours | None | 3 hours |
| *Flash freeze* | None | None | Before lysis | None |

The other major discriminant between common extract preparation protocols is in the post-lysis processing, consisting of the run-off reaction, dialysis, and centrifugation steps. Both the run-off and dialysis steps originate from the pioneering work of Nirenberg and Matthaei [5] and Zubay [4]. During the run-off reaction, the extract is incubated at 37^°^C to allow ribosomes to “run off” their native transcripts, freeing them to translate recombinant transcripts [5, 64]. The clarified extract is then dialyzed against buffer in a step hypothesized to remove unwanted metabolic byproducts that accumulate during the run-off reaction such as inorganic phosphate [7, 56, 59]. Previous work showed that protein synthesis from a T7 expression cassette is not improved in BL21-sourced extracts by including the run-off reaction or dialysis [55, 65]. However, we hypothesized that these steps might impact protein synthesis under control of native bacterial promoter systems. We thus aimed to systematically study the impact of these post-lysis processing steps on protein expression from the native *E. coli* RNAP.

As an initial test, we directly adapted our clarification protocol to incorporate the classic run-off and dialysis steps (Figure 1A, “Post-lysis processing”, green line). Consistent with previous work, in the same reaction conditions, we observed no difference in T7-driven protein synthesis from the additional clarification steps (1220 ± 85 ng/μL post-lysis processed vs. 1240 ± 89 ng/μL non-processed). However, we observed a five-fold improvement in yield from the *E. coli* σ^70^ reporter containing the RNA stability hairpin to greater than 300 ng/μL (Figure 1B and 1C green bar). Overall, these results supported our goal to design a CFE platform that permits robust expression from the native *E. coli* RNAP machinery. However, these experiments did not shed light on why CFE expression under control of bacterial but not T7 promoters is so sensitive to the run-off and dialysis steps.

### Yield improvements to cell-free gene expression are independent of genetic part strength

We next sought to understand the improvement in cell-free expression yields from the post-lysis processing steps. Specifically, we hypothesized that because the run-off reaction and dialysis steps had minimal effect on T7-driven expression, these steps selectively improve transcription by the native bacterial polymerase and likely do not affect the translational machinery. We therefore predicted that the post-lysis processing steps would improve protein synthesis for constructs that had a range of native promoter and ribosome binding site strengths.

To test this hypothesis, we measured cell-free expression protein yield from reporter constructs that contained synthetic *E. coli* σ^70^ promoters and ribosome binding sites (RBSs) that were predicted to have a range of strengths. In each case, we maintained the stability hairpin in the 5’ untranslated region. Protein yield was then quantified using eight-hour kinetic CFE experiments at the 10 μL scale in a 384-well plate to enable continuous monitoring of sfGFP production.

For all σ^70^ reporter designs tested, comparison of reactions using these extracts revealed at least a three-fold improvement in final protein yield from the post-lysis processed extract relative to the non-processed extract (Figures 2A and B). The improvement was most dramatic for the weakest predicted RBS and promoter designs, for which no expression of sfGFP could be observed over background in the non-processed condition. This result is important for two reasons. First, it shows that the enhanced protein yield from the *E. coli* transcriptional machinery due to post-lysis processing is general regardless of regulatory element strength, suggesting that the omission of these steps causes transcriptional inhibition. Second, these data underscore the importance of post-lysis processing for cell-free expression from reporter constructs that have weak native expression—for example, gene circuits or biosensors that are designed with a low ON state to prevent leak.

**Figure 2.**
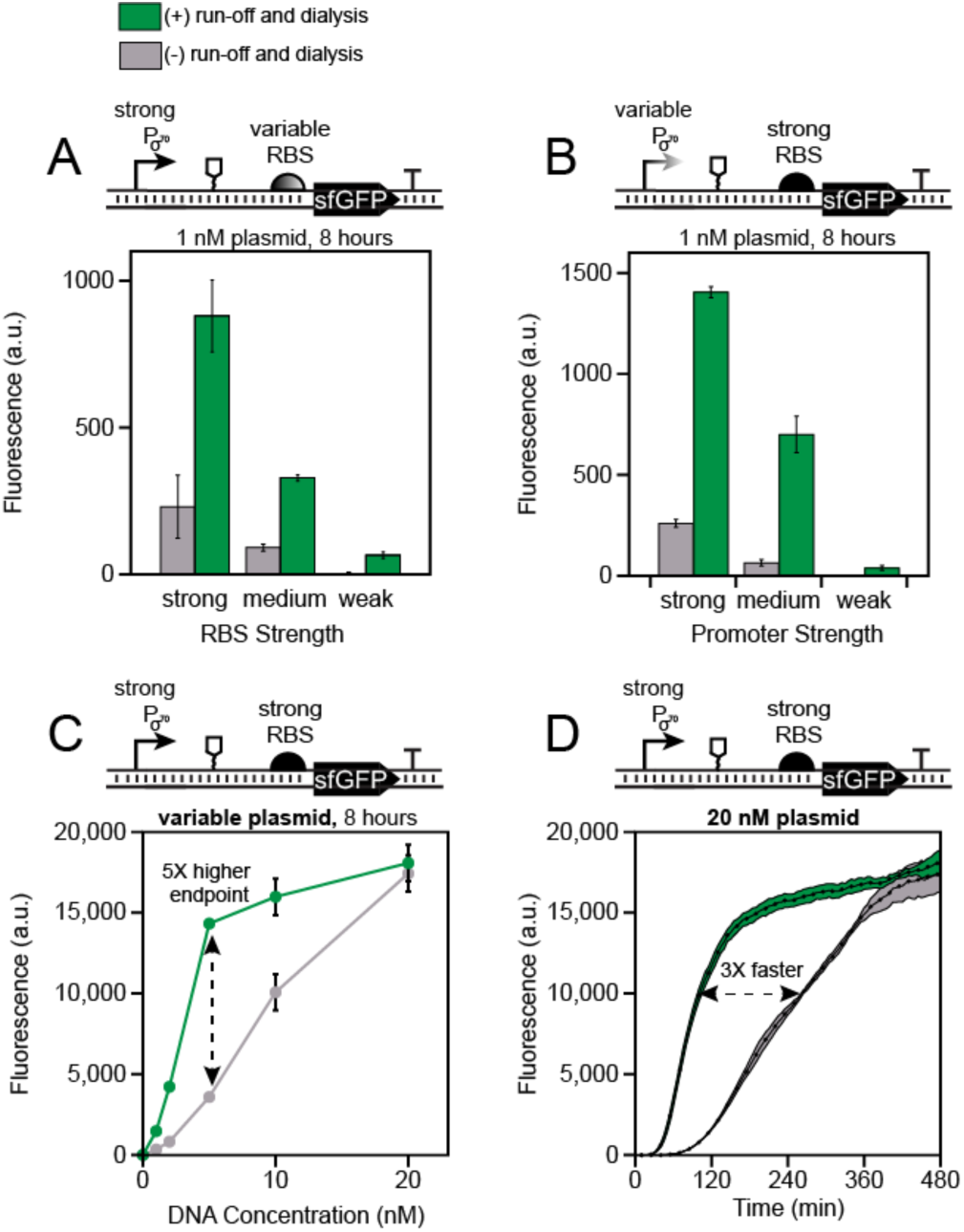
Post-lysis processing steps in extract preparation enhance cell-free gene expression yields from *E. coli* σ^70^ promoters. Extracts prepared with run-off and dialysis (green) increases the yield of an sfGFP reporter across constructs containing a range of synthetic (a) ribosome binding site and (b) *E. coli* σ^70^ promoter strengths, compared to extracts prepared without these steps (gray). (C) Titration of reporter construct DNA that contains a strong promoter and RBS in both extracts suggests that protein synthesis saturates above 10 nM in the processed extract but continues to increase in the non-processed extract. (D) Kinetics at the 20 nM reporter template DNA concentration show that despite having equal endpoint sfGFP production levels, protein production is 3X faster for the processed extract. Endpoint data in (a-c) are from eight-hour experiments incubated and measured in a plate reader at 30^°^C. Error bars represent the standard deviation of the mean across three technical replicates drawn from a single batch of extract.

Based on these observations, we hypothesized that if transcription by the bacterial polymerase is rate-limiting for gene expression in cell-free conditions, then protein production should scale directly with the DNA template concentration until either the available polymerase is saturated or transcription is no longer limiting. To test this, we performed kinetic CFE reactions as above on the strongest promoter and RBS construct over a range of template DNA concentrations (Figure 2C). As predicted, although the processed extract showed marked improvement over the non-processed extract at low DNA concentrations, when 20 nM DNA template was supplemented, both reactions saturated at the same final protein levels (Figure 2C), demonstrating that adding more template DNA can relieve transcriptional limitations. Interestingly, however, the two different extracts varied greatly in the kinetics of protein synthesis (Figure 2D), with the processed extract reaching its endpoint value about three times faster than the non-processed extract. Indeed, the quicker onset of sfGFP production in the processed extract was observed across all template concentrations (Supplementary Figure S2).

We observed a much weaker dependence of CFE yield on DNA template concentration when the sfGFP reporter gene was transcribed by the T7 RNAP. Specifically, increasing the template concentration from 1 nM to 5 nM improved expression from the bacteriophage promoter by 17% in the non-processed and 49% in the processed extract (Figure S3). By contrast, when expression was driven by the endogenous bacterial promoter, final protein yield increased about 10-fold in both processed and non-processed extracts (Figure 2C).

### Yield improvements to cell-free gene expression arise from enhanced transcriptional activity

Taken together, the above results suggest that transcription by the *E. coli* RNA polymerase can be the rate-determining step of a CFE reaction, and that although transcriptional constraints can be partially relaxed by adding more DNA template, the transcription activity *in vitro* can be linked to specific steps of the extract preparation. To validate our hypothesis that transcription limits cell-free gene expression, we next aimed to remove the confounding effect of reporter protein translation and quantify the transcription rate in each extract by monitoring gene expression at the RNA level. To do this, we replaced the sfGFP coding sequence in our strongest σ^70^ and T7 reporter template plasmids with the malachite green aptamer (MGA) sequence. Once transcribed, this RNA aptamer binds to the malachite green fluorophore and causes the dye to fluoresce. The MGA system therefore enables a convenient fluorescence-based quantification of transcript levels and, as such, has been previously used both in cells and *in vitro* [66-68]. As expected, under the consensus *E. coli* σ^70^ promoter, we observed substantially higher expression of the MGA reporter in the processed extract over the course of a three-hour experiment (Figure 3A).

**Figure 3.**
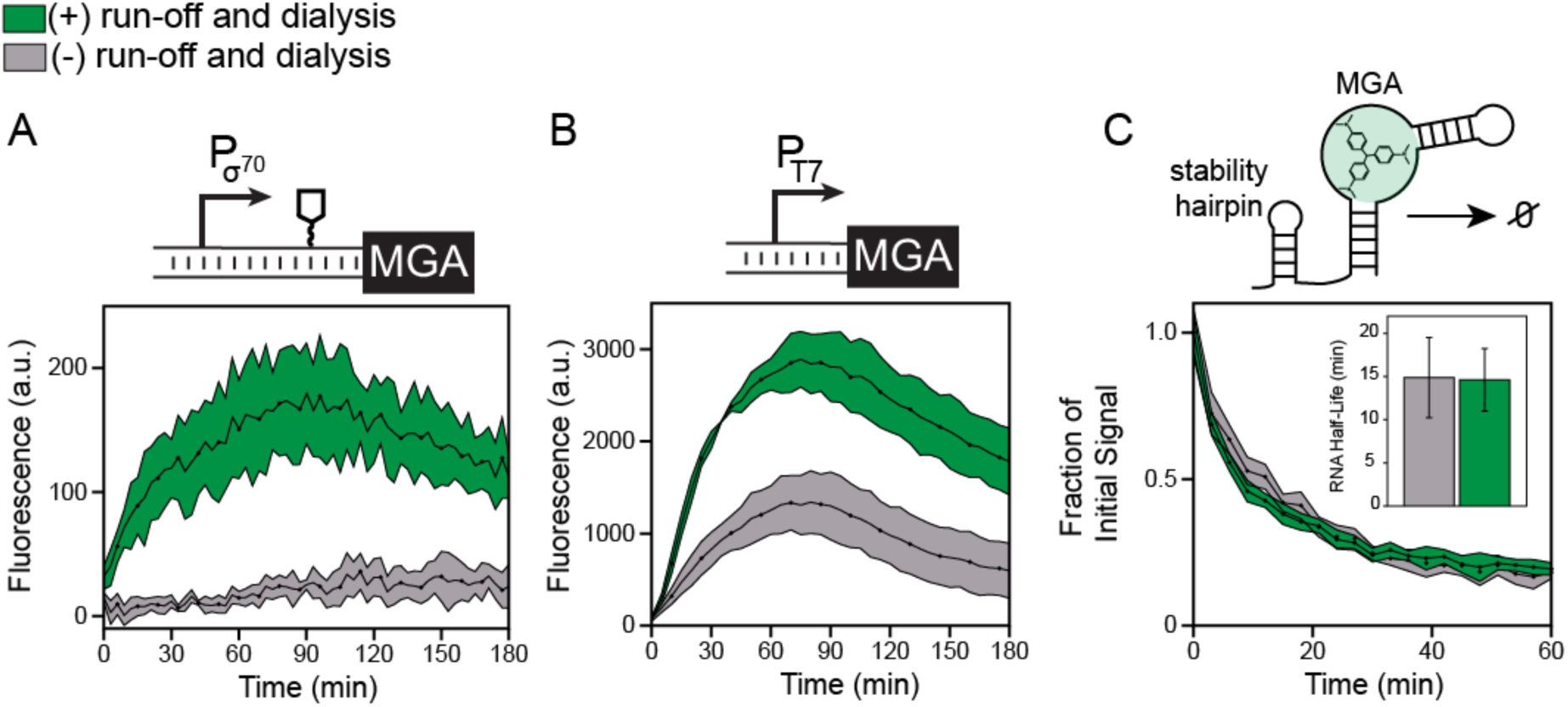
Post-lysis processing enhances the transcription rate in CFE systems. (a) *In vitro* transcription of the malachite green RNA aptamer (MGA) from a strong *E. coli* σ^70^ promoter in a cell-free gene expression reaction containing the malachite green dye using either processed (green) or non-processed (gray) extracts. The construct encoded a 5’ stability hairpin before the MGA DNA sequence to enhance fluorescence observation. (b) *In vitro* transcription of the MGA construct from a T7 promoter and no 5’ stability hairpin. Kinetic data in (a) and (b) represent the average of three technical replicates, with shaded region representing plus-minus one standard deviation of the mean. (c) Characterization of RNA degradation rates. The malachite green RNA aptamer was purified with the 5’ stability hairpin, mixed with the malachite green dye and 10% extract by volume and the decay in fluorescence over time was measured to quantify RNA degradation rates (k_degradation_). Inset: the RNA half-life (ln(2)/k_degradation_) was estimated by fitting an exponential decay function to the fluorescence kinetics over the first 30 minutes to fit (k_degradation_). Errors in half lives are propagated from three independent measurements and reported as standard deviation.

Notably, we observed almost no fluorescence from the MGA reporter in the non-processed condition, suggesting that RNA degradation keeps pace or outstrips weaker transcription rates in those extracts. The hypothesis that RNAse-mediated degradation controls transcript abundance is also supported by the observation that there is a peak in MGA expression at ∼80 minutes, likely caused by continued degradation after transcriptional slow-down from the depletion of NTPs in the reaction [69].

Interestingly, when transcribed from the T7 promoter, MGA production is also about two-fold higher in the processed extract (Figure 3B). This result contradicts our earlier finding that the post-lysis processing steps do not appreciably improve expression from the phage polymerase (Figure 1B). However, these results can be explained together by the presence of translational bottlenecks in CFE that limit protein but not mRNA production from the bacteriophage promoter - only by removing the translational bottlenecks can we directly observe differences between extract preparation strategies for the T7 reporter. Thus, the extract processing steps enhance *in vitro* transcription generally, regardless of the polymerase, but are especially important in transcriptionally limiting conditions, especially the expression of proteins from less productive bacterial transcriptional machinery.

Rather than activating transcription, an alternative mechanism for how the post-lysis processing steps improve extract productivity would be the stabilization of transcripts by removing or inhibiting RNAses present in the extract. In fact, batch-to-batch variation in extract productivity has previously been observed to be partly due to variability in RNA degradation rates between batches [35]. To test for this in our extract protocol variations, we measured RNA degradation by spiking purified MGA and the malachite green dye into both processed and non-processed extracts and observed the decay of fluorescence over time [35] (Figure 3C). Interestingly, we observed nearly identical half-lives for the RNA in each extract, indicating that the post-lysis processing steps do not alter RNA degradation rates. Additionally, similar measurements performed with MGA RNAs lacking a 5’ stability hairpin showed that the hairpin increases the RNA half-life by about two-fold, underscoring the importance of RNA stability in transcriptionally-limited systems (Supplementary Figure S4).

Overall, these experiments prove that the improvements in CFE yield from post-lysis processing are due to improvements in transcriptional output of the reactions. Importantly, this effect is enhanced in reactions where mRNA levels are limiting.

### Protocol sensitivity can be assessed in transcriptionally-limiting conditions

Having demonstrated that transcriptional limitations in CFE reactions can be lifted by post-lysis processing through a run-off reaction and dialysis, we next aimed to measure the sensitivity of extract productivity to other extract preparation protocol variations. Specifically, we sought to measure the impact that three common protocol variations would have on CFE reactions operating under transcriptional limitations. These include media conditions used to grow cells prior to lysis, flash freeze steps used as a convenient stopping point between growth and lysis, and the individual impacts of run-off and dialysis (Figure 4). From a practical standpoint, this analysis would facilitate the adoption of extract-making protocols to new labs with less experience with the nuances of preparing extract. Moreover, the sensitivities could reveal useful insight into how each extract preparation protocol step impacts the overall productivity of the extract.

**Figure 4.**
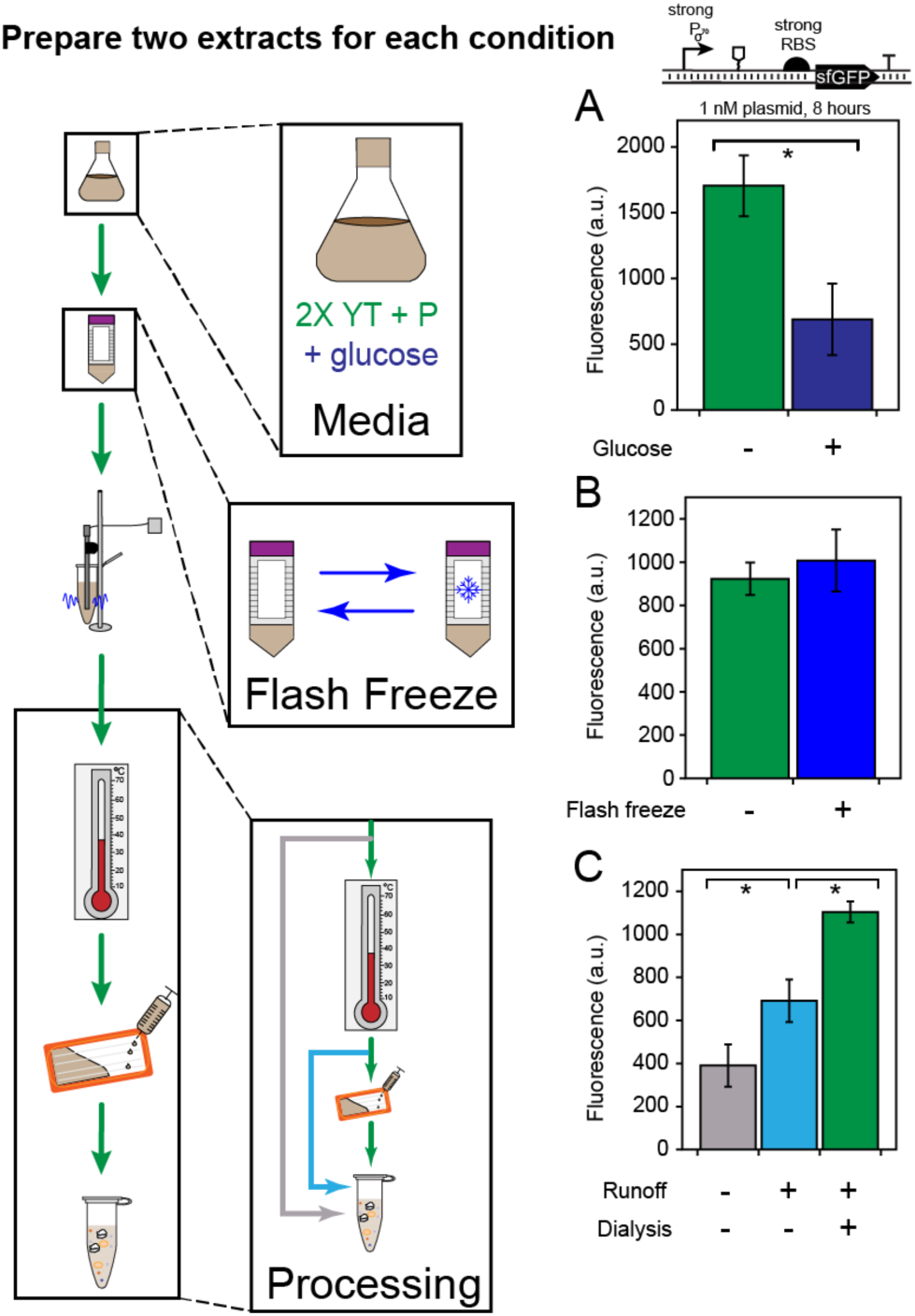
Extract activity is sensitive to some protocol variants. (a) Addition of glucose (navy) to the 2X YT + P culture media reduces CFE yields from bacterial promoters under transcriptionally limiting conditions (p = 0.035). (b) Flash freezing the cell pellet before lysis does not significantly impact the extract’s productivity (p = 0.30). (c) Extract clarification by the Kwon et. al. [55]protocol for BL21 (gray), with run-off reaction (light blue), and with run-off reaction and dialysis (green) under transcriptionally-limiting conditions shows that both the run-off and dialysis steps contribute to the observed increase in transcriptional activity (p = 0.039 for green vs. gray and p = 0.031 for green vs. light blue). All bars represent the average of six independent cell-free reactions, drawn from three technical replicates of two independently generated extracts made on different days. Error bars represent the standard deviations of the six measurements. In each case, unpaired one-tailed Student’s t tests were performed for N = 2 extracts using Welch’s correction for heteroscedasticity.

Before starting this analysis, we first had to assess the impact of batch-batch variability on our ability to discriminate meaningful differences between extract preparation protocol variations. Previous work has shown that yields from bacterial promoters in transcription-translation reactions can vary by more than 50% between batches intended to be identical [34]. To estimate batch-batch variability in our base protocol, we prepared three identical extracts on three different days, aiming to replicate all experimental conditions in the preparation. We then performed identical, transcriptionally-limited sfGFP synthesis reactions on all three extracts on three different days, with aim of determining whether observed variation was due to extract-extract variation, experiment-experiment variation, or both. Our results (Supplementary Figure S5) showed that both extract and experimental variation are statistically significant (p = 0.030 for extract variability and 0.026 for experimental variability from a two-factor ANOVA). Despite these differences, the rankings for both extract and experimental productivity showed high correlation across all three days. Moreover, the combination of those two conditions was the only one found to be statistically significantly different than any other experiment-extract combination in pairwise t-tests.

Compared against previous studies, we observed lower batch-batch variability between individual extracts; using the endpoint fluorescence averaged across all three days, the coefficient of variation (CV) between the three extracts was 11%. We hypothesize that one reason for this consistency was the use of sonication for cellular lysis rather than less precise techniques like bead-beating since sonication allows the amount of energy delivered to the sample to be precisely controlled. Nevertheless, achieving precise reproducibility between extract preparations remains difficult. Despite preparing all media and buffers in tandem and inoculating the starter cultures from a clonal population, we still observed differences in growth conditions such that the time required to reach the target harvest optical density varied from 3.5 to 4.5 hours. This is one possible explanation for the observed batch-batch variability.

We also observed a small amount of experiment-experiment variability (CV = 11%), which was just as much as extract-extract variability. We attribute this variability to small variations in pipetting plasmid DNA from a concentrated stock into the reaction master mix, given the previous observation that protein synthesis yields under transcriptional limitations are exquisitely sensitive to template DNA concentration (Figure 2C). Because this amount of experimental variability will be the same for every reaction measured in a given experiment, we felt confident in our ability to discriminate extract productivities with statistically significant differences in final protein titers.

Having established our ability to discriminate extract batch variability from meaningful protocol differences in extract preparation, we next aimed to measure our protocol’s sensitivity against specific variations. We tested variation at every step except the sonication, which was kept constant as a design constraint, since previous work has shown that the sonication energy has a strong, strain-dependent impact on extract productivity [55]. Specifically, the three variables that we tested were media composition for cell growth, flash freezing prior to lysis, and the post-lysis processing steps individually rather than as a group, paying close attention to major protocol variations that exist in the literature with the aim of creating a robust protocol. In each case, we performed the same sfGFP synthesis reactions under limiting bacterial transcription for 8 hours, but also prepared two independent batches of extract following each protocol variant to remove any unknown bias present in single batches. Under these conditions, we expected to see the maximum sensitivity between protocol variants.

First, we set out to test different media formulations. Two common formulations used in the literature include 2X YT + P [56] and 2X YT + P + G [55], where P stands for buffered potassium phosphate and G is for glucose. Although our earlier extracts were prepared with 2X YT + P media, we sought to test if glucose could be supplemented to the media and bacterial transcription activity retained. We therefore tested the impact of growing lysate chassis cells in media with and without glucose on the extract’s performance. Surprisingly, supplementation of glucose significantly diminished our observed yields under bacterial transcription control (Figure 4A, p = 0.035). While glucose was initially implemented by Kim and Choi [70] to lower phosphatase activity in the lysate (i.e., hexose phosphatase), and maintained in the protocols developed by Jewett and Swartz to activate natural energy metabolism [16, 17, 71], its presence in the media is deleterious to endogenous transcription in the lysate. Thus, growing the cells on 2x YT + P was selected as a core design parameter moving forward.

Next, we aimed to test the impact of flash-freezing cells in liquid nitrogen prior to lysis. In the literature, flash-freezing has been an optional step implemented in some but not all protocols and has a practical advantage as a convenient stopping point if included. We therefore wondered if flash freezing would diminish the quality of the extract for expression from bacterial promoters. Encouragingly, we observed that flash-freezing our cell pellet immediately before lysis has no appreciable impact on the final cell-free yield (Figure 4B). Since the additional clarification steps contribute five hours of labor to the extract protocol, the ability to split the procedure across more than one day would greatly broaden its appeal, particularly for laboratories acclimating to the methods for extract preparation.

Finally, we sought to examine the impact of each post-lysis clarification step individually, particularly the result of performing a run-off reaction incubation but not dialysis. We observed 50% recovery in the yield from the bacterial promoter from the run-off reaction with no dialysis, suggesting that the two steps have specific yet different effects on the physiochemical makeup of the extract (Figure 4C).

### A wide array of transcriptional genetic circuits are functional in a modified cell-free extract preparation framework

Having observed the relaxation of transcriptional constraints in the constitutive expression of a reporter protein off a bacterial promoter, we next aimed to demonstrate the activation of a number of synthetic gene expression circuits driven by bacterial genetic machinery in our modified CFE system. We assembled a library of gene expression activators and repressors, functioning at both the RNA and protein levels, with a diversity of expression levels and tested them in both processed and non-processed extracts (Figure 5). The six circuits conditions that we assayed were: small transcription-activating RNAs (STARs) [72]; RNA transcriptional repressors [73]; the inducible protein LuxR transcription factor [74]; CRISPR-Cas9-mediated DNA cleavage [75, 76]; toehold switch translation activating RNAs [77]; and a translational theophylline riboswitch [78]. The individual reaction conditions for each circuit are presented in Supplementary Table 1. In each case, the reaction’s ON state was first maximized by adjusting magnesium concentration, a crucial determinant of CFE functionality, and then adjusting the concentration ratios for each plasmid supplied to the reaction.

**Figure 5.**
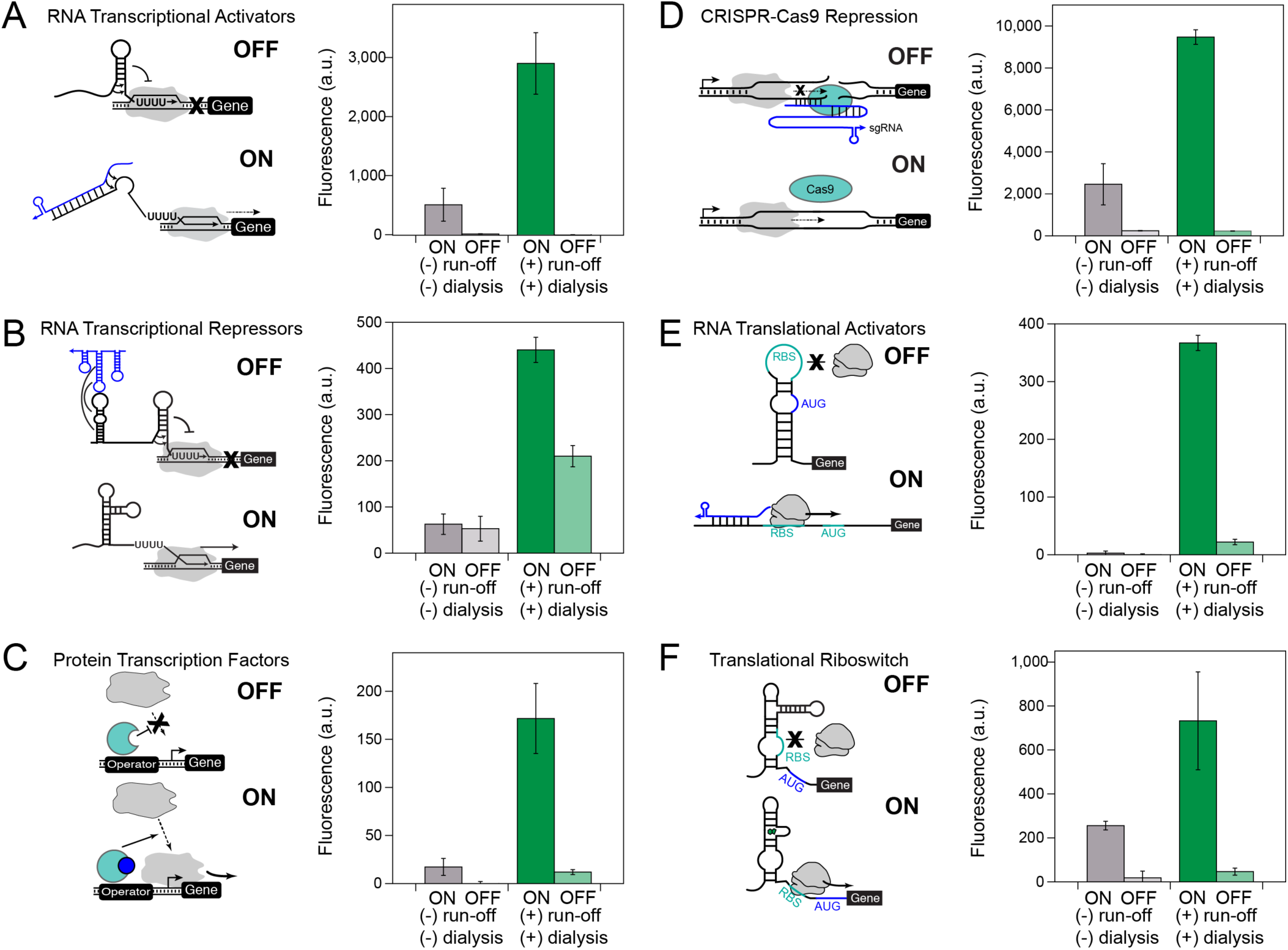
Processed extracts enable cell-free expression of a wide array of genetic circuitry. Post-lysis processing enables much higher ON states, including activation in some cases, for (a) small RNA transcriptional activators (STARs), (b) RNA transcriptional repressors, (c) LuxR, the N-acyl homoserine lactone (AHL)-inducible σ^70^ transcription factor, (d) CRISPR-Cas9 cleavage, (e) toehold switch RNA translation activators, and (f) translational riboswitches. Each experiment was done in technical triplicate in either a post-lysis processed (green) or non-processed (gray) extract for 8 hours at 30^°^C in a plate reader monitoring sfGFP production. The total plasmid DNA concentration was held constant in each reaction between the ON and OFF states of each circuit to remove variation caused by unwanted cell-free transcription and translation of the antibiotic resistance genes encoded on each plasmid. Complete reaction conditions including plasmid and inducer concentrations and optimal magnesium levels are included in Supplementary Table 1. Kinetics of gene expression trajectories for each circuit are presented in Supplementary Figure S6.

In all six circuits that we tested, the ON state for the processed extract was at least three times higher than the ON state for the non-clarified extract. In two circuits (transcription factors, toeholds), we could not observe signal above background in the non-processed extract. For the transcriptional repressors, the ON state for the non-processed extract was nearly indistinguishable from the OFF state (15% repression vs. 52% repression in the optimal extract). Although the circuits designed to have greater ON activity were at least functional in the non-processed extract, they were much slower: the STARs, CRISPR-Cas9, and riboswitch systems all responded around three times faster in the optimized extract (see Supplementary Figure S6 for all kinetics). Since speed of response is a major advantage for cell-free biosensors over more conventional diagnostic methods, our extract system could to play an important role in the proliferation of this technology. Taken together, it is clear that post-lysis processing is a critical step to generate a functioning extract system that is broadly applicable to diverse genetic circuits.

## Summary

In this work, we established a new framework for production of cell-free extracts for *in vitro* synthetic biology. With this modified procedure, we demonstrated ∼3-5-fold increases in cell-free production of a model reporter placed under control of a bacterial promoter. Equally important, we identified transcription from the σ^70^ promoter to be a key limiting step in CFE operating at non-saturating plasmid DNA concentrations. This result opposes a common prevailing notion, mainly borne from experiments with bacteriophage polymerases, that energy regeneration for translation strictly determines final protein titer in *in vitro* transcription-translation reactions [79-81]. With our optimized extracts, we were able to activate a diverse array of genetic circuitry in CFE, including RNA transcriptional activators, RNA translational activators, protein transcription factors, CRISPR-Cas9 nuclease activity, RNA translational repressors, and a translational riboswitch.

We observed that the key process improvements for activating cell-free gene expression from bacterial promoters were the inclusion of an 80-minute run-off reaction and subsequent dialysis. We hypothesize that dialysis removes a small molecule global inhibitor of transcription that accumulates in the crude lysate during the run-off reaction. However, due to the complex and largely unknown metabolome of the extract, we were unable to identify the origin of this inhibition. Future studies may do well to further elucidate the mechanism, with an ultimate aim toward engineering a better source strain.

Although improved expression of proteins from bacterial promoters in cell-free extracts has been shown previously, previous work has been insufficient to characterize what exact process steps are necessary to activate bacterial transcription. This lack of understanding, compounded by disparities in reaction conditions and reagent compositions between published methods, makes it exceedingly challenging for new labs to adopt cell-free gene expression technologies. Additionally, as CFE systems gain popularity for prototyping genetic regulators, especially in non-model organisms, the link between extract preparation and resulting transcriptional activity will grow in importance. By illuminating the importance of extract processing steps and highlighting their relevance to CFE, we anticipate that our work will expand the utility of cell-free systems throughout synthetic biology.

## MATERIALS AND METHODS

### Plasmid Construction and Purification

A table of all plasmids and the DNA concentration used in each experiment of this study can be found in Supplementary Table 1, and plasmid architectures can be found in Supplementary Data File 1. Plasmids were assembled using either Gibson assembly or inverse PCR and purified using a Qiagen QIAfilter Midiprep Kit. pJBL002 and pJBL004 are from [82] and pJBL7022 is based on pJBL006 from[82]. pJBL3859, 4860, 5816, and 5817 are from [83]. pJBL7008 is based on the theophylline riboswitch RS.E from [78]. pJBL2812 and 2814 are from [39]. pJBL7023 is from [55]. pJBL623 and 632 were a gift from Dr. Lei Qi.

### Extract Preparation

Cell-free extract was prepared based on the protocol in [55]. Rosetta 2^TM^ cells were streaked onto chloramphenicol agar plates overnight, then inoculated into 30 mL LB supplemented with antibiotic and grown in a 250 mL baffled flask at 37^°^C. After 16 hours, 20 mL of the stationary culture was used to inoculate 1L of autoclaved growth media 2X YT+P (16 g/L tryptone, 10 g/L yeast extract, 5 g/L sodium chloride, 7 g/L potassium phosphate dibasic, 3 g/L potassium phosphate monobasic) in a 2.5L baffled Tunair flask. A separate bottle containing 7.2% glucose solution was autoclaved and added at the time of inoculation for 2X YTP+G experiments to a final glucose concentration of 1.8%.

Cultures were grown to the exponential phase optical density 3.0 ± 0.2 for approximately 4 hours at 37^°^C and 200 rpm. The culture was divided in two and centrifuged for 15 minutes at 5,000xg at 4^°^C to pellet the cells. The cell pellets were washed three times in 25 mL Buffer A (50 mM Tris, 14 mM Mg-glutamate, 60 mM K-glutamate, 2 mM DTT, brought to pH 7.7 with acetic acid) and re-centrifuged at 5,000xg at 4^°^C for 10 minutes. After the fourth and final centrifugation at 7,000xg at 4^°^C for 10 minutes, pellets were re-suspended in 1 mL Buffer A/g pellet and transferred to 1.6 mL micro-centrifuge tubes. For the flash freeze experiments, cell pellets were submersed in liquid nitrogen before this step and thawed the next day.

The cell suspensions were then lysed by sonication on a Q125 sonicator with a 3.175 mm diameter probe at a frequency of 20 kHz and 50% amplitude by 10 second ON/OFF pulses for a total of 60 seconds (delivering ∼300J). The lysate was centrifuged for 10 minutes at 4^°^C and 12,000xg, and the supernatant was removed. For preparations including a run-off reaction, the tubes of crude lysate were covered in aluminum foil and were incubated shaking with lids on for 80 minutes at 37^°^C and 200 rpm. The extract was re-centrifuged for 10 minutes at 4^°^C and 12,000xg, and the supernatant was removed. For preparations including a dialysis, the extract was injected into a 10K MWCO Slide-a-Lyzer dialysis cassette (ThermoFisher, Catalog #66380) and dialyzed against 600 mL of Buffer B (5 mM Tris, 14 mM Mg-glutamate, 60 mM K-glutamate, 1 mM DTT, pH 8.2) at 4^°^C for three hours. The dialyzed extract was removed from the cassettes and centrifuged once more for 10 minutes at 4^°^C and 12,000xg. The supernatant was removed, aliquoted, and snap-frozen in liquid nitrogen.

### CFE Experiment

The final CFE reaction mixture is composed of the following reagents: 10-20 mM magnesium glutamate; 10 mM ammonium glutamate; 130 mM potassium glutamate; 1.2 mM ATP; 0.850 mM each of GTP, UTP, and CTP; 0.034 mg/mL folinic acid; 0.171 mg/mL yeast tRNA; 2 mM amino acids; 30 mM PEP; 0.33 mM NAD; 0.27 mM CoA; 4 mM oxalic acid; 1 mM putrescine; 1.5 mM spermidine; 57 mM HEPES; 30% CFE extract by volume; plasmid DNA to the desired concentration (refer to Supplementary Table 1 for plasmid DNA concentrations used in this study); and water. For reactions involving T7 RNAP expression, in-house purified T7 RNAP was doped into the reaction at 0.10 mg/mL. The optimal magnesium concentration was determined for each reporter construct and was found to be 16 mM magnesium glutamate in nearly all cases (for exceptions to this, as well as the inducer concentrations in Figure 5, refer to Supplementary Table 1).

All kinetic CFE reactions were prepared on ice in triplicate at the 10 μL scale. 33 μL of a mixture containing the desired reaction components was prepared and then 10 μL was pipetted into three wells of a 384-well plate (Corning, 3712), avoiding bubbles. Plates were sealed (ThermoScientific, 232701) and sfGFP fluorescence (emission/excitation: 485/520 nm; gain = 50) was monitored every five minutes on a BioTek Synergy H1m plate reader for 8 hours at 30^°^C. For the bulk endpoint experiments in Figure 1, reactions were prepared to the 15 μL scale in triplicate, pipetted into the bottom of a 2.0 mL micro-centrifuge tube, and incubated without shaking at 30^°^C overnight for 15 hours. Final sfGFP titers were calculated from a previously developed plate reader correlation. For the malachite green experiments in Figure 3, fluorescence (emission/excitation: 615/650; gain = 100) was measured every three minutes. For all experiments, a no-DNA negative control was prepared in triplicate for every extract being tested. All reported fluorescence values have been baseline-subtracted by the no-DNA condition, and all error bars include the propagated error from the no-DNA condition.

### RNA Purification and RNA Degradation Experiments

The malachite green RNA aptamer with and without the 5’ stability hairpin was purified from a runoff *in vitro* transcription of pJBL7004 or pJBL7005 using in-house prepared T7 RNAP. The RNA was purified by denaturing urea-polyacrylamide gel electrophoresis on an 8% polyacrylamide gel, followed by elution into nuclease-free water overnight and ethanol precipitation.

For RNA degradation experiments, 2.2 μM of the purified RNA was doped into a 11 μl mixture containing 10% malachite green dye and 10% extract by volume, with the balance nuclease free water, quickly pipetted into a 384-well plate, and malachite green fluorescence (emission/excitation: 615/650; gain = 100) was monitored every three minutes for two hours. Half-lives were then calculated by fitting the fluorescence trajectories over the first half hour to the equation F = F_0_e^-kt^, where t_1/2_ = ln(2)/k.

## Acknowledgments

We extend a special thanks to our collaborators for thoughtful insight into and editing of this manuscript. This work was supported by the Air Force Research Laboratory Center of Excellence for Advanced Bioprogrammable Nanomaterials (C-ABN) Grant FA8650-15-2-5518 (to M.C.J.), the U.S. Defense Advanced Research Projects Agency’s (DARPA) Living Foundries program award HR0011-15-C-0084, the David and Lucile Packard Foundation (to M.C.J.), an NSF CAREER Award (1452441 to J.B.L.), the Camille Dreyfus Teacher-Scholar Program (to M.C.J. and J.B.L.), and Searle Funds at the Chicago Community Trust (to J.B.L.). A.D.S. was supported in part by the National Institutes of Health Training Grant (T32GM008449) through Northwestern University’s Biotechnology Training Program. The US Government is authorized to reproduce and distribute reprints for Governmental purposes notwithstanding any copyright notation thereon. The views and conclusions contained herein are those of the authors and should not be interpreted as necessarily representing the official policies or endorsements, either expressed or implied, of the Air Force Research Laboratory, Air Force Office of Scientific Research, DARPA, or US Government.

